# Rapid experience-dependent tuning of spinal and transcortical stretch reflexes supports goal-directed movement

**DOI:** 10.64898/2026.04.29.721632

**Authors:** Taiyeba Akter, Robin Rohlén, Per Petersson, Michael Dimitriou

**Affiliations:** Department of Medical and Translational Biology, Umeå University, Umeå, Sweden; Department of Diagnostics and Intervention, Umeå University, Umeå, Sweden; Department of Bioengineering, Imperial College London, London, UK; Department of Experimental Medical Science, Lund University, Lund, Sweden

## Abstract

The short-latency stretch reflex (SLR) is the fastest sensorimotor response in human limbs. The spinal SLR scales with background muscle activity and is affected by recent movement history. But adaptive plasticity of spinal reflexes has classically been studied through operant conditioning, which induces an arbitrary change in reflex size over days to weeks. In contrast, rapid, experience-dependent feedback adaptation is often attributed to transcortical mechanisms underlying the long-latency reflex (LLR). Whether task constraints can progressively shape the SLR within a single session remains unclear. Here, naive human adults (n = 16; 9 females, 7 males) performed an instructed-delay reaching task while holding a robotic manipulandum which delivered brief haptic perturbations during action preparation. We recorded high-density surface electromyography (64-channel arrays) from the pectoralis major and posterior deltoid to investigate reflex gains. Pre-perturbation muscle activity showed no systematic goal-directed change. With a long-enough preparatory delay (750 ms), SLR gains decreased progressively with experience when the planned movement was expected to stretch the homonymous muscle, rather than shorten it. This tuning occurred in both muscles, was evident within individual participants, and was accompanied by a corresponding decrease in reflexively generated force opposing perturbations toward the cued target. No such tuning was detected with a short preparatory delay (250 ms). LLR responses were goal-directed from the first trial blocks and progressed in a comparable manner across the session, at both preparatory delays. Our study shows that spinal feedback circuits can progressively adapt within a single session to support the performance of goal-directed movements.

**Significance statement:** Reflexes mediated by spinal circuits are often regarded as fixed, automatic responses, with rapid experience-dependent adjustment reserved for longer-latency reflex pathways that route through the brain. Plasticity of the fastest, spinal stretch reflex has instead been associated with days to weeks of dedicated training. Here we show that, as healthy adults repeatedly perform a reaching task within a single session, the spinal stretch reflex is progressively tuned according to the planned movement, reducing resistance to motion toward the goal. Rapid adaptation of feedback control is therefore not confined to neural pathways traversing the brain but extends to spinal circuits, highlighting that the entire reflex loop contributes to the progressive optimization of goal-directed behavior.

## Introduction

Voluntary goal-directed movements are prepared before execution. During this preparatory phase, neural activity evolves toward a state that supports the forthcoming action before any overt movement occurs (Cohen et al., 2010; Haith and Bestmann, 2018). Studies using the instructed-delay paradigm, in which a target is cued in advance of a Go cue that triggers movement, have revealed widespread goal-directed preparatory activity across premotor and primary motor cortices, as well as subcortical pathways (Prut and Fetz, 1999; Cisek and Kalaska, 2005; Churchland et al., 2006a; Churchland et al., 2006b).

A prominent conceptual framework proposes that preparatory activity establishes an internal dynamical configuration (sets ‘the initial conditions’ of a neural population) rather than issuing explicit commands to muscles (Churchland et al., 2010; Shenoy et al., 2013). This framework helps reconcile evidence of robust preparatory activity in central circuits with the repeated observation that overt muscle activation shows little to no consistent goal-directed modulation during preparation, at least when measured with conventional single-channel electromyography (EMG). However, how preparation manifests as improved motor performance remains unclear.

One emerging view is that preparation also involves tuning the sensitivity of proprioceptive feedback circuits. Muscle spindles, the sensory or ‘afferent’ limb of the stretch reflex, can be tuned in a goal-directed manner during movement preparation, with relative inhibition of spindle responses from muscles expected to stretch during the forthcoming reach (Papaioannou and Dimitriou, 2021). Such tuning offers a mechanism for proactively shaping motor feedback behavior (i.e., reducing antagonist stiffness, thereby facilitating its stretch during movement) without requiring overt changes in muscle activity before movement. Accordingly, a categorical or ‘set-related’ decrease of stretch reflex gains has been repeatedly shown in muscles about to be stretched (Papaioannou and Dimitriou, 2021; Torell et al., 2023a, b; Rohlén et al., 2025), including at short (spinal) latencies, traditionally considered “too early” to be flexibly configured (Scott, 2016).

The short-latency stretch reflex (SLR) is mediated by spinal circuits, and represents the fastest sensorimotor response in human limbs. The spinal SLR scales with baseline muscle activity i.e., “automatic gain-scaling” (Pruszynski et al., 2009), can depend on limb configuration (Weiler et al., 2019), as well as recent stretch history (Stratmann et al., 2024). However, adaptive plasticity of spinal reflexes has classically been studied through operant conditioning, which induces an arbitrary change in reflex size that develops over days to weeks (Wolpaw et al., 1983; Thompson and Wolpaw, 2014). By contrast, the long-latency (transcortical) stretch reflex (LLR) is widely regarded as task-dependent and adaptable on an iterative basis (e.g., Pruszynski and Scott, 2012; Cluff and Scott, 2013; Coltman and Gribble, 2020). These findings have fostered the general belief that rapid, experience-dependent tuning is a feature of long-loop reflex pathways that route through the brain, and not of spinal feedback circuits. Whether task constraints can progressively shape the SLR within a single session remains unclear. Such tuning is plausible, as SLR gain may be adjusted at several points along the reflex arc, including at the muscle spindles, via independent fusimotor control (Dimitriou, 2021b, a, 2022).

Here, we investigate whether the SLR shows progressive, experience-dependent modulation in naive participants performing an instructed-delay reaching task without prior practice. We recorded high-density surface EMG (HDsEMG; 64-channel arrays) from the pectoralis major and posterior deltoid to quantify reflex responses with improved robustness and spatial sampling, while a robotic manipulandum delivered brief kinematic perturbations during movement preparation. We expected progressive SLR inhibition when participants prepared to stretch the homonymous muscle (congruent target), but not when preparing to shorten it (incongruent target), and only with a sufficiently long preparatory delay. We also examined whether this tuning had mechanical consequences at the hand, and whether the LLR progressed in the same manner. We show that spinal reflex gains can undergo rapid, experience-dependent tuning within a single session, in a manner typical of transcortical circuits.

## Materials and Methods

### Participants

A total of 16 right-handed and neurologically healthy individuals (age: 28.6 ± 4.6 years; 9 females, 7 males) participated in this study. The participants were naive as to the purpose of the study and performed no practice trials before data collection began. All were financially compensated and gave informed written consent before participating. The study was conducted in accordance with the Declaration of Helsinki and was approved by the Swedish Ethical Review Authority (reference number: 2025-01785-02).

### Experimental setup

#### Robotic platform

The participants sat upright in a customized adjustable chair with a supported forehead in front of the Kinarm robotic platform (Kinarm End-Point Robot, BKIN Technologies). They used their right hand to grasp the handle of the robotic manipulandum (Fig. 1A). The right forearm was placed inside a customized foam structure, resting on an airsled that allowed frictionless arm movement in a 2D plane. To ensure a secure mechanical connection, the forearm, hand, Kinarm handle, and foam-cushioned airsled were secured using a leather fabric with Velcro attachments. This attachment also kept the wrist in straight alignment with the forearm throughout the experiment, while the left hand was placed parallel to the right hand at a safe distance and rested on a soft cotton towel. The robotic platform measured the hand’s position, and sensors inside the robotic handle recorded the forces exerted by the participants’ right hand (six-axis force transducer; Mini40-R, ATI Industrial Automation). Position and force data were sampled at 1 kHz.

**Figure 1.**
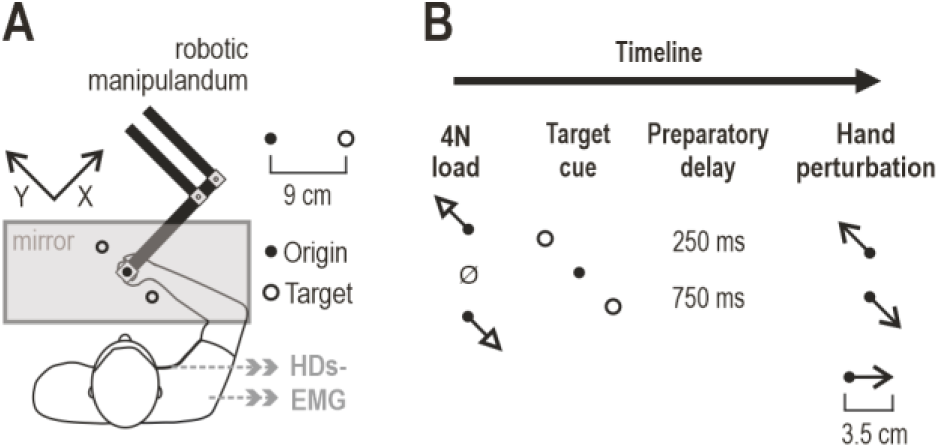
Experimental setup and instructed-delay reaching task. ***A***, Participants (n = 16) held the graspable end of a robotic manipulandum with their right hand. Vision was directed at a one-way mirror displaying a projected monitor image, and hand position was represented by a cursor. The right forearm rested on an air sled and the hand was immobilized at the wrist. Two targets were displayed 9 cm from the central origin, in the +Y (135°) and −Y (315°) directions. A 64-channel HDsEMG array (4 mm inter-electrode distance) recorded pectoralis activity (8 × 8 array), and another such array recorded posterior deltoid activity (13 × 5 array). ***B***, Each trial began by slowly loading the hand to 4 N in the upper-left (+Y) or lower-right (−Y) direction, or there was no load (Ø; “null”). During this phase, participants maintained the hand at the origin. One of two visual targets (+Y or −Y) was then cued (turned red) for a short (250 ms) or long (750 ms) preparatory delay, after which the hand was perturbed 3.5 cm in the +Y or −Y direction (toward or away from the cued target) over 150 ms, while cursor position remained frozen at the origin. At perturbation end, the Go signal occurred (target turned green), and participants moved promptly to the target.

#### High-density surface electromyography

Two 64-channel HDsEMG electrode arrays (8 × 8 and 13 × 5 arrays, the latter with one corner electrode absent; 4 mm inter-electrode distance, OT Bioelettronica, Torino, Italy) were placed on the skin surface of the pectoralis major and posterior deltoid muscles, respectively. The arrays were placed with their columns parallel to the muscle fiber direction, as estimated by palpation. Before placing the two HDsEMG arrays, the skin surface was prepared by shaving (if necessary) and cleaned with abrasive paste to reduce impedance. A double-sided adhesive foam was then attached to the electrodes, and the electrode cavities were filled with conductive gel to facilitate signal conductivity. The HDsEMG arrays were secured with surgical tape to promote good electrode contact with the skin throughout the session. The respective wireless amplifiers (Muovi+, OT Bioelettronica, Torino, Italy) were also secured parallel to them with tape. A ground electrode (Dermatrode HE-R Reference Electrode type 00200–3400, 5 cm diameter, American Imex) was placed on the processus spinosus of the seventh cervical region.

The HDsEMG amplifiers transmitted their data wirelessly to the SyncStation+ (OT Bioelettronica, Torino, Italy), where the EMG signals were recorded in a monopolar configuration at 2000 Hz, amplified, 16-bit A/D-converted, with a 150x gain, and bandpass-filtered between 10 and 500 Hz. Brief occasional losses of wireless transmission appeared as lags between an amplifier’s data stream and the trigger record; each lag was corrected by realigning the subsequent signal to the trigger record, with the lost samples set to zero so that the affected trials could be identified and excluded offline (see ‘Data processing’).

#### Synchronization of recording systems

To achieve temporal alignment between kinematic and electrophysiological recordings, the Kinarm robotic system and the HDsEMG acquisition were synchronized offline in a two-stage procedure. First, the Muovi+ amplifiers had internal trigger pulses that were aligned with the internal trigger pulse of the SyncStation+. Second, to synchronize the Kinarm system with the SyncStation+, it generated a rectangular pulse via one of its analogue output channels at the start and end of each trial. This signal was recorded through an auxiliary input channel of the SyncStation+. These synchronization pulses were subsequently used to align the Kinarm-derived data with the HDsEMG recordings offline. The accuracy of this alignment is on the order of 1 ms (synch pulses sampled at 2kHz; 0.5 ms between data points), ensuring accurate temporal correspondence across all data streams.

### Experimental task

The experimental design of this study used a delayed-reach paradigm, incorporating systematic variations in target direction, muscle load, preparatory delay and perturbation direction (Fig. 1B). A one-way mirror prevented direct visual feedback of the hand and robotic handle, while visual stimuli were projected onto the mirror in the plane of movement. These stimuli included a 1 cm diameter white cursor that moved in a one-to-one correspondence with the hand’s position (cursor position was frozen during haptic perturbations). Additionally, two targets were continuously displayed as orange circular outlines, positioned 9 cm from the central starting point. The targets were located in the front-left and back-right directions, corresponding to 135 degrees (‘+Y’) and 315 degrees (‘−Y’), respectively (see Fig. 1). Reaching along this axis has previously been shown to preferentially engage the pectoralis major and posterior deltoid (Papaioannou and Dimitriou, 2021).

Each participant completed a total of 20 blocks of trials, where each block consisted of 24 unique trials presented in a block-randomized order, and comprising three load directions, two cue targets, two preparatory delays and two perturbation directions (3 × 2 × 2 × 2 = 24 trial conditions; 480 trials in total). Specifically, each trial began by moving the cursor to an “origin” circle in the middle of the workspace (Fig. 1A). For the trial to progress, the cursor had to remain immobile inside the origin for a random wait period between 1 and 1.5 s, after which a 4 N load (800 ms rise time and 1200 ms hold time) was applied in either the +Y or −Y direction, or there was no load (‘null’ load). One of the targets was then cued by suddenly turning from an orange outline to a filled red circle. After a pre-defined preparatory delay (250 or 750 ms), a position-controlled perturbation of the hand was applied (3.5 cm displacement, 150 ms of rise time, no hold period), swiftly moving the hand in the +Y or −Y direction. Each haptic perturbation was programmed to reproduce the kinematics of a fast, naturalistic point-to-point movement, approximating a bell-shaped velocity profile. To ensure the target kinematics were achieved on every trial, the robot was permitted to engage its maximum available stiffness (∼40,000 N/m) as needed. At the perturbation offset, the ‘Go’ cue appeared (the cued target turned green), and participants had to reach that target promptly, as previously instructed.

A trial was considered complete once the participant held their hand stationary within the target for 300 ms, at which point visual feedback on performance was delivered. Trials were classified as ‘Correct’ if the target was reached within 1.4 s of the Go cue onset, and as ‘Too slow’ otherwise. A correct trial incremented the block score recorded by the task program. Following performance feedback, a new trial began when the cursor was returned to the origin. Since trials were presented in a block-randomized order, the direction of perturbations was unpredictable to participants, even after experiencing a particular load, target cue and preparatory delay. Participants were given a one-minute rest after every three consecutive blocks of trials, up to the eighteenth block (the additional two blocks were executed without a scheduled break). Additionally, participants were allowed to take short unscheduled breaks if they wished, by moving the cursor to the side of the workspace. Trials in which the perturbation moved the hand toward the cued target are termed ‘congruent’, and trials in which it moved the hand away from the cued target are termed ‘incongruent’. Analyses focused on the 12 conditions in which the perturbation stretched the homonymous muscle (i.e., −Y perturbations for the pectoralis and +Y perturbations for the posterior deltoid).

### Data processing

The raw monopolar HDsEMG signals were band-pass filtered between 20 and 500 Hz (third-order zero-phase Butterworth filter) to remove movement artifacts, and notch filtered at the power-line frequency (50 Hz). From these signals, we computed the difference between adjacent channels along the muscle fiber direction to attenuate far-field contributions and improve spatial selectivity, reducing inter-channel crosstalk prior to signal aggregation. This procedure reduced the number of signals from 64 to 56 for the pectoralis muscle (8 × 8 array) and from 64 to 59 for the posterior deltoid (13 × 5 array). The HDsEMG signals, already band-limited to 500 Hz, were then down-sampled from 2 kHz to 1 kHz (every second sample retained) to match the Kinarm data. The time-series data of each trial was then epoched from 200 ms before to 800 ms after perturbation onset (defined below). Artifact rejection was performed at the level of individual channels: on each trial, a channel was excluded if its amplitude (median absolute value over the trial epoch) deviated from the median amplitude of all channels on that trial by more than 4 robust standard deviations (1.4826 × median absolute deviation across channels). On average, 10.0% of pectoralis and 7.5% of posterior deltoid channels were excluded per trial, leaving approximately 50 and 55 channels, respectively, for aggregation. Then, each remaining channel was full-wave rectified and, at each sample, the median across channels was computed, giving one EMG signal per muscle and trial. The median was chosen as the aggregation statistic due to its robustness to few channels having very small or very large amplitudes. The resulting inter-channel temporal offsets between adjacent electrodes (∼1 ms at the inter-electrode spacing used, given a conduction velocity of 4 m/s) are negligible compared with the temporal resolution relevant for our stretch reflex analyses (≥ 25 ms).

As each block value rests on a single trial per condition, the reliability of the aggregated EMG was assessed by split-half analysis across channels. For each participant, the retained channels were divided at random into two halves, block values were computed from each half as from the full grid, and the two sets were correlated (Pearson) across blocks, separately for congruent and incongruent trials. Correlations were averaged on the Fisher z scale over the two congruencies and 50 random splits, and Spearman-Brown corrected. Reliability was high for both muscles, both reflex epochs (defined below) and both preparatory delays (group medians 0.98–0.99; individual values 0.91–1.00; Figure S1).

At the level of whole trials, trials without recorded EMG were removed (i.e., classified as ‘missing data’), separately for each muscle. These were trials in which the EMG signal was absent for the entire trial or contained a transmission gap within the analyzed window (−50 to 100 ms relative to perturbation onset; signal below 0.1% of the participant’s typical median amplitude for at least 10 consecutive ms). Missing data resulted from technical issues during recording, principally losses of wireless transmission, an unexpectedly depleted amplifier battery in the final two blocks of one participant, and trials for which the EMG could not be matched to the Kinarm trial record. These technical issues occurred in episodes that typically spanned one or more consecutive blocks of an individual participant, affecting all conditions of a block equally. Further, the posterior deltoid recording of one participant showed a contact fault during the final 13 trials of the session (abrupt bursts of pre-perturbation activity locked to the perturbation direction, with loss of the stretch response, confined to the deltoid array); these trials were also removed. In total, 5.2% of pectoralis and 7% of posterior deltoid trials in the analyzed conditions were removed. Described across participants, 5.3 ± 6.1% (mean ± SD) of pectoralis and 7.0 ± 6.8% of posterior deltoid trials were removed. As all contrasts were conducted within participants, from blocks with available data, missing EMG data only reduced the number of trials the participant’s estimates were based on. Because these missing data episodes affected all conditions of a block equally, missing data were balanced across conditions and congruencies (Figure S2). Trials without EMG were excluded from EMG analyses only i.e., their handle force and kinematic data were retained. In addition, 43 trials of one participant were not recorded by the Kinarm system and were therefore absent from all analyses.

EMG signals were z-normalized separately for each participant and muscle, enabling comparison across participants, in line with previous work (e.g., Dimitriou, 2018; Torell et al., 2023a; Rohlén et al., 2025). The mean and standard deviation were computed from the aggregated EMG signal (median across rectified channels), pooled across all samples of the epoch and all trials of all 24 conditions (missing-data trials excluded). This normalization yields results equivalent to normalization relative to the maximal trial-averaged value (Torell et al., 2023b). All trials were time-aligned to movement onset (hereafter ‘perturbation onset’). Perturbation onset was defined on each trial as the time at which hand velocity reached 5% of peak velocity observed during the position-controlled perturbation, a criterion commonly used to define movement onset (e.g., Pigeon et al., 2003; Sainburg and Schaefer, 2004). Position and force data were rotated by 45 degrees so that the y-axis of the rotated frame lay along the axis of targets, loads and perturbations (i.e., task axis; Fig. 1A). Position and force data were low-pass filtered at 20 Hz (third-order zero-phase Butterworth filter), and force was signed so that positive values opposed the perturbation and, for analyses, was baseline-adjusted on the 50 ms preceding perturbation onset. Force was not normalized.

The principal statistical analyses focused on conditions in which the muscle was mechanically stretched by the perturbation. All relevant trials were analyzed, regardless of reaching success. As each block contained one trial per condition, EMG was averaged (mean) over the samples of each epoch to give one value per participant, condition and block. As the position-controlled perturbations were kinematically matched across conditions, differences in reflex magnitude between conditions are indicative of differences in reflex gain. Accordingly, goal-directed tuning of reflex gain was assessed as the difference in EMG activity between congruent and incongruent trials. Reflex responses were analyzed in two epochs that follow the conventional division of the SLR (25–50 ms post-perturbation onset) and LLR (50–100 ms), in line with previous work with both arm and shoulder muscles (e.g., Weiler et al., 2015; Scott, 2016; Rohlén et al., 2025). Background (pre-perturbation) activity was quantified in the 50 ms epoch preceding perturbation onset. Because reflex onset differs slightly between individuals due to limb length and other factors (Schuurmans et al., 2009), and 20 ms has also been used at times as the onset of the SLR for the posterior deltoid (e.g., Kurtzer et al., 2016), we verified the onset of the SLR and LLR epochs in our data. A two-segment function adapted from Weiler et al. (2015), comprising a constant level followed by a linear rise, with the breakpoint constrained to ≥ 15 ms, was fitted to each participant’s stretch-trial EMG of the posterior deltoid (750 ms preparatory delay, all loads, blocks 11–20). For this, each participant’s stretch trials (both congruencies) were averaged and baseline-corrected (pre-perturbation epoch), and the function was fitted by least squares (breakpoint in 1-ms steps) from 5 ms until the smoothed response (5-ms moving average) first reached half its peak within 20–70 ms. The response had begun by 25 ms in 11 of 16 participants (onset 22.7 ± 6.1 ms, mean ± SD; 95% CI 19.5–25.9 ms). The 50 ms boundary between the SLR and LLR was also supported by the congruent-trial EMG, which did not begin to decline consistently until 58 ms (95% bootstrap CI 45–64 ms; Figure S3), i.e., after the end of the SLR epoch. This onset was defined on the group-mean congruent response (each participant scaled to its peak stretch response within 20–99 ms; 3-ms moving average) as the start of the run, within 35–65 ms, in which the slope was at least half its steepest negative value; the CI was obtained from 1000 bootstrap resamples of participants. A similar onset of the congruent decline was observed for the pectoralis (52 ms, 95% bootstrap CI 43– 53 ms). Throughout, each sample is taken to represent the 1 ms beginning at it, so that consecutive epochs do not overlap. That is, for analyses, the SLR comprises the 25 samples from 25 to 49 ms, and the LLR the 50 samples from 50 to 99 ms. All data preprocessing was performed in MATLAB (Version R2019a, MathWorks).

### Experimental design and statistical analyses

A repeated-measures design was used, and all inference was at the level of participants. For all paired and one-sample comparisons, normality of the differences was assessed with the Shapiro-Wilk test, followed by a *t*-test or, when normality was rejected (*p* < 0.05), a Wilcoxon signed-rank test. Normality was tested with swtest (MATLAB File Exchange). Confidence intervals of means were t-based. Effect sizes are reported as Cohen’s *d_z_* (*t*/√*n*) or, for Wilcoxon tests, as the matched-pairs rank-biserial correlation. All tests were two-tailed.

The correction for multiple comparisons followed the structure of our hypotheses. Previous afferent and reflex findings (Papaioannou and Dimitriou, 2021; Torell et al., 2023b, a; Rohlén et al., 2025) led us to expect goal-directed tuning of the SLR only when the preparatory delay is sufficiently long (above ∼300 ms), and the homonymous muscle is about to be stretched. On this basis, all other cases, such as those involving short preparatory delays or preparation for muscle shortening, could be regarded as controls rather than as tests of the same hypothesis. However, these earlier findings were obtained with single-channel electrodes, whereas the multi-channel recordings used here may detect modulation that single electrodes miss, and our short preparatory delay (250 ms) is close to the minimum delay established with single-channel methods. We therefore included the short preparatory delay in the same family of hypotheses: for each muscle and reflex epoch, the primary test was performed at both preparatory delays and Bonferroni-corrected for the two tests (α = 0.025). No additional correction was applied across the two muscles, which provide independent replications of the same hypothesis. The SLR was the primary epoch of interest. The LLR was analyzed to assess whether it progressed across the session in the same manner as the SLR; for this epoch, group curves across blocks are shown for descriptive purposes, and the primary test was the participant-level correlation analysis described below.

Reaching performance was quantified as the percentage of successful trials per block and congruency, pooled across participants, for the conditions in which the analyzed muscle was stretched by the perturbation, at each preparatory delay. The curves were described with an exponential, y = A + B·exp(−block/τ), fitted by least squares and drawn when significantly different from a constant (F-test, p < 0.05). For inference, each participant’s success rate in the ‘late’ trial blocks (11–20) minus that in the ‘early’ blocks (1–10) was tested against zero, separately for each congruency. Performance in the two congruencies was not compared with each other, as the stretched muscle has a different role in each.

For each participant, congruency, epoch and block, EMG was averaged (mean) across the three load conditions. Collapsing across loads simplifies the analysis to a single contrast per muscle, epoch and preparatory delay, avoiding a threefold multiplication of tests. It also asks whether goal-directed tuning occurs regardless of the pre-perturbation mechanical state of the muscle, because an effect that is evident after averaging across unloading, null and preloading conditions is unlikely to depend on a particular level of background activity. As congruent and incongruent trials occurred equally often with each load, collapsing does not confound congruency with load. The load conditions are nevertheless not expected to be strictly equivalent, as reflex gain may scale with background activity (Pruszynski et al., 2009). Where relevant, analyses also examined the no-load condition in isolation, i.e., responses in the absence of any external (pre-) load (see below). Although group curves across blocks are shown for descriptive purposes, the primary test for the SLR was fit-free: for each participant and congruency, the mean of the late blocks (11–20) minus the mean of the early blocks (1–10) was computed, and this change was compared between congruent and incongruent trials with a paired test. The change within each congruency is shown along with the 95% confidence interval. The same analysis was applied to the pre-perturbation epoch as a control for congruency-specific changes in background activity across the session; this comparison was evaluated at α = 0.05 without correction, i.e., the more conservative choice for detecting a potential confound.

Progression of reflex tuning (SLR and LLR) across blocks was also characterized within each participant. This analysis was performed with all three load conditions and, separately, with the no-load condition alone, for both muscles. For each participant, congruency and epoch, block values were averaged across the selected load conditions, and the Spearman rank correlation (ρ) between EMG and block number was computed over the blocks with data, as an index of monotonic change that makes no assumption about the shape of the progression. To remove any contribution of background activity, the partial rank correlation of reflex EMG with block, controlling for pre-perturbation EMG, was also computed. For this, the block values of all selected loads were pooled and rank-transformed, the mean of each load condition was removed from the ranks of reflex EMG, pre-perturbation EMG and block number (load as a fixed effect), and the partial correlation was computed on the pooled residuals, so that background and reflex activity were related within each load condition. With the no-load condition examined in isolation, the load term drops out and pre-perturbation EMG was partialled out within participant only. The coefficients were Fisher *z*-transformed and compared between congruent and incongruent trials with a paired test; plotted values are shown back-transformed with their 95% confidence intervals. For the SLR, both partialled and unpartialled coefficients are plotted. Because the LLR is known to be less sensitive to gain-scaling, unpartialled values are plotted, but the relevant partialled values are also reported in the Results section. The same correlation computed on pre-perturbation EMG served as a control for background activity. For the SLR, this analysis characterizes how the change identified by the primary test progresses within participants, and it was not included in the correction for multiple comparisons; for the LLR, the unpartialled correlation constituted the primary test and was corrected as described above.

Handle force along the main task axis was analyzed to establish whether the change in SLR EMG had mechanical consequences at the level of the whole limb. Force also provides the link between reflex tuning and behavioral improvement: a decrease in reflex-based resistance to hand displacement toward the target would be expected to facilitate reaching it. The SLR epoch was shifted by 30 ms for both muscles (i.e., force epoch 55–80 ms after perturbation onset), a value at the lower end of the electromechanical delays reported for upper-limb muscles during voluntary contractions (Norman and Komi, 1979) and goal-directed reaching (Keyser et al., 2019). Force was taken from all trials of the stretch conditions, including those with missing EMG, and averaged across loads as for the EMG. Mean force in this epoch was analyzed with the same late-minus-early contrast and paired comparison between congruent and incongruent trials. As force is recorded in the same trials as the EMG (and additionally in trials without EMG), it corroborates rather than independently tests the reflex result, and it was therefore not included in the correction for multiple comparisons. All statistical analyses were performed in MATLAB (Version R2019a, MathWorks); the custom code used for statistical analyses and the relevant data are available at [repository, DOI - tbd].

## Results

In this study, 16 participants performed the classic instructed-delay reaching task while holding a robotic manipulandum with their right hand. We simultaneously recorded hand kinematics, handle forces, and HDsEMG signals from the right pectoralis major and posterior deltoid. In each trial, the hand was perturbed toward or away from the cued target after which the participants had to promptly complete the reach to the cued target. A single experimental session comprised 20 blocks of trials, each containing one repetition of all 24 conditions in randomized order. Analyses focused on trials in which the perturbation stretched the muscle of interest. In “congruent” trials, the perturbation was in the direction of the cued target, whereas in “incongruent” trials the perturbation was in the direction opposite to the target.

Figure 2*A* shows the responses of an exemplary participant in trials where the pectoralis was stretched following a long preparatory delay (late trial blocks, pooled across load conditions). Despite identical displacements during the perturbation, there is a clear goal-directed difference in the pectoralis SLR (25–50 ms): the response is weaker when preparing to stretch the pectoralis (congruent target; blue) rather than shorten it (incongruent target; purple). This difference became more pronounced in the LLR epoch (50–100 ms), whereas pre-perturbation activity was similar across target conditions. Figure 2*B* shows the spatial distribution of the pre-perturbation and reflex activity aggregated in Figure 2*A*. Reflex activity varied across recording sites, and in this participant the goal-directed difference was distributed across much of the grid rather than confined to a particular region. A recording grid covering a large part of the muscle surface is likely to capture reflex activation more comprehensively than a single-channel electrode at one site.

**Figure 2.**
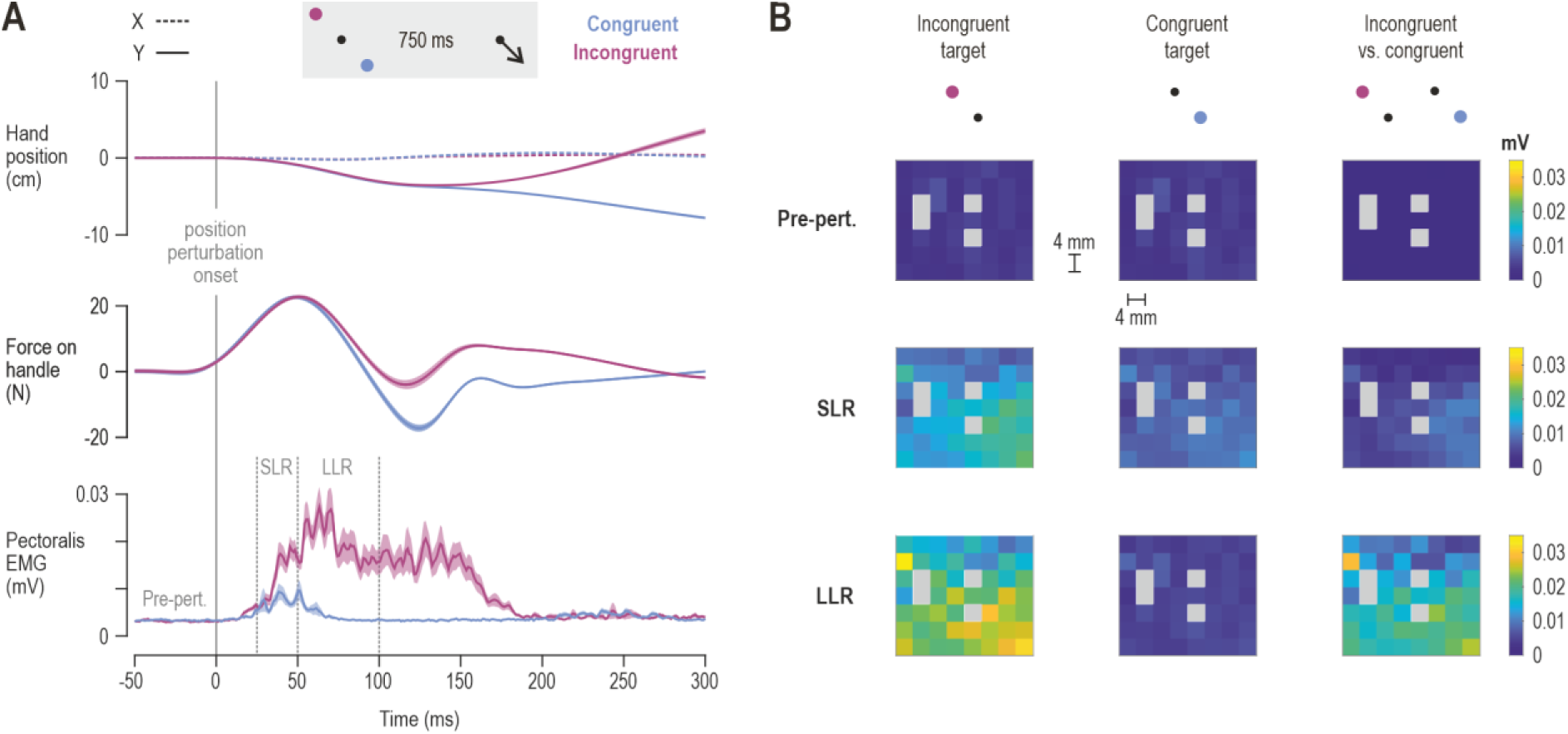
Exemplary single-participant responses. ***A***, Data from one participant in trials with a long preparatory delay (750 ms) and a perturbation stretching the pectoralis (−Y direction), averaged across the three load conditions. The cued target was either in the +Y direction (incongruent target; purple) or −Y direction (congruent target; blue). Traces show the mean across the late trial blocks (11–20; 30 trials each for incongruent and congruent targets, 10 per load condition) and shading represents ±1 SEM. Time zero denotes perturbation onset. Top: hand position along the X (dashed) and Y (solid) axes. The negligible displacement along the X axis indicates that movement was confined to the task axis (Y). Middle: force on the handle along the Y axis. Bottom: averaged pectoralis EMG (mV). Vertical dash-dotted lines delimit the short-latency stretch reflex (SLR, 25–50 ms) and long-latency stretch reflex (LLR, 50–100 ms) epochs; the pre-perturbation epoch (‘pre-pert.’) spans the 50 ms window prior to perturbation onset. Despite matched perturbation kinematics, EMG activity already differs in the SLR epoch as a function of the cued target. ***B***, Spatial distribution of pectoralis activity across the HDsEMG grid in the pre-perturbation, SLR and LLR epochs, for the trials shown in *A*, with incongruent (left) and congruent (middle) targets. Each cell represents one single-differential channel (4 mm inter-electrode distance), drawn at its upper electrode; its value is the full-wave rectified signal averaged over the epoch and then across trials (mV). The right column shows the difference between incongruent and congruent trials (negative differences shown as zero). Gray cells denote channels excluded by channel-level artifact rejection (see Materials and Methods).

### Goal-directed SLR tuning with a long preparatory delay

A trial was classified as successful if the participant reached the cued target within a predefined time window after the Go signal (see Materials and Methods). With a long preparatory delay (750 ms), success rate increased across trial blocks for both congruent and incongruent targets, whether the planned movement would stretch the posterior deltoid (Fig. 3*A*) or the pectoralis (Fig. 3*E*), approaching a plateau in the second half of the session. The progression was well described by an exponential function (posterior deltoid, congruent: *R*^2^ = 0.86, *p* = 4.6 × 10^−8^; incongruent: *R*^2^ = 0.84, *p* = 2.2 × 10⁻⁷; pectoralis, congruent: *R*^2^ = 0.93, *p* = 8.3 × 10⁻¹¹; incongruent: *R*^2^ = 0.9, *p* = 2.2 × 10⁻⁹). Paired tests indicated that, compared with early trial blocks (blocks 1–10), success rate was significantly higher in late trial blocks (blocks 11–20) in all cases (posterior deltoid, congruent: *t*(15) = 3.11, *p* = 0.0072, *d*z = 0.777; incongruent: *t*(15) = 6.2, *p* = 1.7 × 10⁻⁵, *d*z = 1.55; pectoralis, congruent: *t*(15) = 4.51, *p* = 0.0004, *d*z = 1.127; incongruent: *t*(15) = 5.2, *p* = 0.0001, *d*z = 1.3).

**Figure 3.**
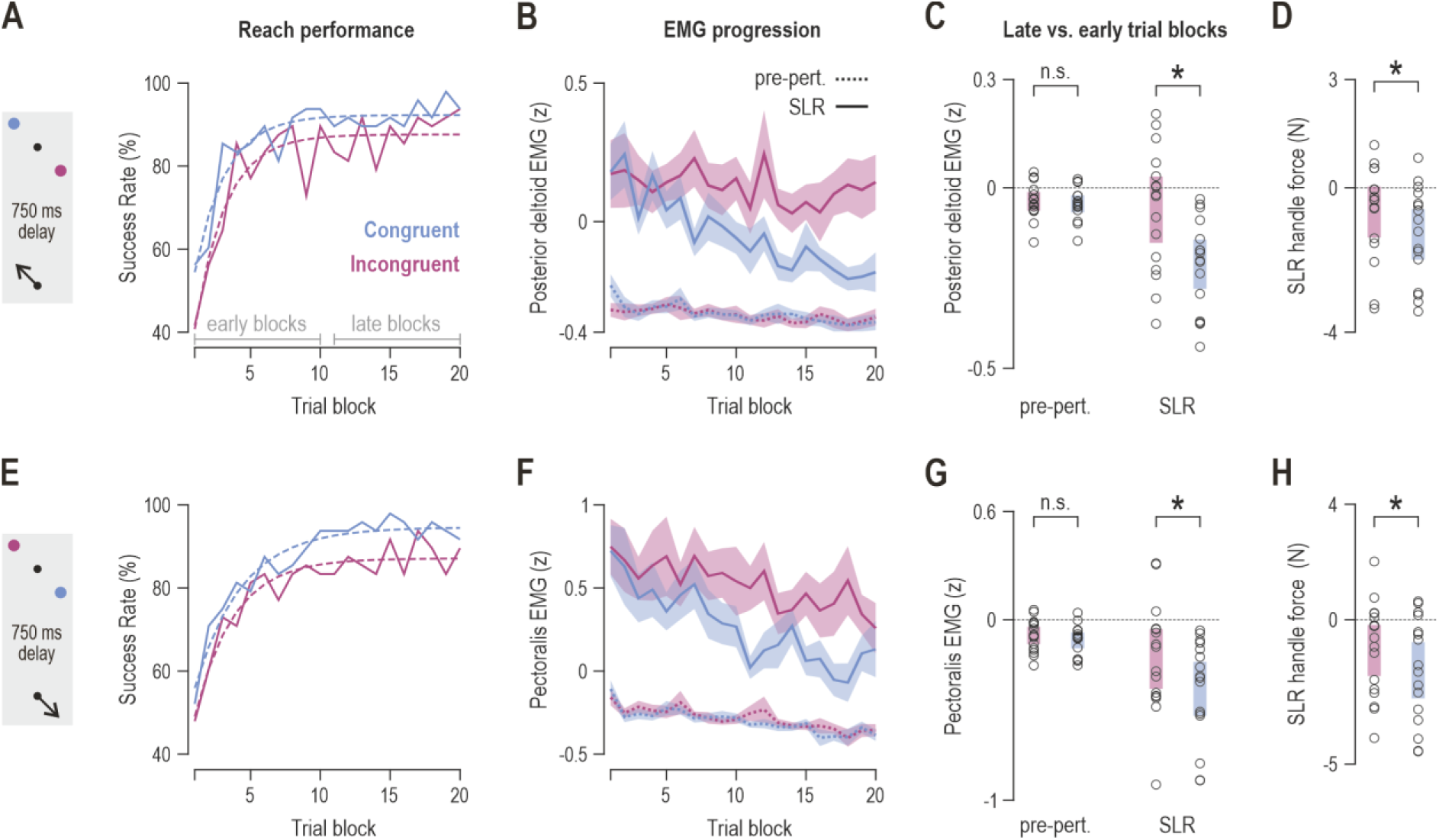
Goal-directed SLR tuning develops over the session with a long preparatory delay. All data are from trials with a relatively long preparatory delay (750 ms) and a perturbation stretching the respective muscle (schematics on the left; n = 16). ***A***, Reaching success rate across trial blocks for congruent (blue) and incongruent (purple) targets, that is, when preparing to stretch or shorten the posterior deltoid, respectively. Dashed lines show descriptive exponential fits. Gray brackets indicate the early (1–10) and late (11–20) trial blocks. ***B***, Mean z-scored posterior deltoid EMG across trial blocks in the pre-perturbation (dotted lines) and SLR epochs (solid lines). Shading represents ±1 SEM. Note the progressive decrease of SLR responses when preparing to stretch the muscle (congruent target; blue). ***C***, Change in pre-perturbation and SLR activity from early to late trial blocks (late − early) in incongruent (purple) and congruent (blue) trials. Each circle represents a single participant, and shaded bars represent 95% confidence intervals. ***D***, As *C*, but for force on the handle (task axis Y) in the SLR epoch shifted by a 30 ms electromechanical delay. ***E–H***, As *A–D*, but for the pectoralis major. Single stars indicate statistically significant paired comparisons, at α = 0.025 for SLR comparisons (Bonferroni correction across the two preparatory delays) and at α = 0.05 for all other comparisons (see main text for more details); n.s., not significant.

Reflex responses were quantified as z-scored EMG activity in each epoch (see Materials and Methods). For each participant, change across the session was quantified as the difference in activity between late and early trial blocks (late − early), and this change was compared between congruent and incongruent trials. A more negative change in congruent than in incongruent trials therefore indicates goal-directed inhibition that developed over the session when preparing to stretch the homonymous muscle. In the posterior deltoid, SLR responses decreased across trial blocks when participants prepared to stretch the muscle (congruent target; blue), compared to when preparing to shorten it (incongruent target; purple; Fig. 3*B*). Accordingly, the late-versus-early change in SLR activity was significantly more negative in congruent than in incongruent trials (Fig. 3*C*; *t*(15) = −4.23, *p* = 0.00073, *d*z = −1.057). The pectoralis showed the same pattern (Fig. 3*F*), with SLR responses decreasing more steeply in congruent than in incongruent trials, and a correspondingly more negative late-versus-early change in congruent trials (Fig. 3*G*; *t*(15) = −2.86, *p* = 0.0118, *d*z = −0.715). Pre-perturbation activity decreased slightly across trial blocks, more clearly in the pectoralis, but this decline was common across congruent and incongruent trials (dotted lines in Fig. 3*B*,*F*): late-versus-early change in pre-perturbation activity did not differ between congruencies (posterior deltoid: *t*(15) = −0.86, *p* = 0.4; pectoralis: *t*(15) = −1.23, *p* = 0.24; Fig. 3*C*,*G*).

The SLR tuning was also assessed in the mechanical domain. Force on the handle was quantified within the SLR epoch shifted by an electromechanical delay of 30 ms (see Materials and Methods). For both the posterior deltoid (Fig. 3D; *t*(15) = −2.31, *p* = 0.035, *dz* = −0.576) and the pectoralis (Fig. 3H; *t*(15) = −2.91, *p* = 0.011, *dz* = −0.727), the late-versus-early change in SLR-related handle force was significantly more negative in congruent than in incongruent trials, further indicating that reflexive resistance to perturbations in the direction of the planned movement weakened over the session, which would facilitate reaching the goal.

### No goal-directed SLR tuning with a short preparatory delay

With a short preparatory delay (250 ms; Fig. 4), success rate also improved across the session (Fig. 4*A*,*E*; posterior deltoid, congruent: Wilcoxon *W* = 136, *p* = 0.00047, *r*_rb_ = 1; incongruent: *t*(15) = 10.3, *p* = 3.4 × 10⁻⁸, *d*_z_ = 2.575; pectoralis, congruent: *t*(15) = 4.43, *p* = 0.00048, *d*_z_ = 1.107; incongruent: *t*(15) = 6.04, *p* = 2.2 × 10⁻⁵, *d*_z_ = 1.51). As with the long preparatory delay, the progression was well described by an exponential function (posterior deltoid, congruent: *R*^2^ = 0.93, *p* = 1.5 × 10⁻¹⁰; incongruent: *R*^2^ = 0.87, *p* = 2.5 × 10⁻⁸; pectoralis, congruent: *R*^2^ = 0.87, *p* = 3.6 × 10⁻⁸; incongruent: *R*^2^ = 0.92, *p* = 5.4 × 10⁻¹⁰). However, SLR responses did not diverge substantially between congruent and incongruent trials across trial blocks (Fig. 4*B*,*F*), and the late-versus-early change in SLR activity did not differ between congruent and incongruent trials in either the posterior deltoid (Fig. 4*C*; *t*(15) = −0.66, *p* = 0.522) or the pectoralis (Fig. 4*G*; Wilcoxon *W* = 40, *p* = 0.159). Likewise, there were no significant differences in the late-versus-early change of pre-perturbation activity (posterior deltoid: Wilcoxon *W* = 68, *p* = 1; pectoralis: *t*(15) = 0.25, *p* = 0.81; Fig. 4*C*,*G*) or SLR-related handle force (posterior deltoid: *t*(15) = −0.68, *p* = 0.508; pectoralis: *t*(15) = −0.74, *p* = 0.469; Fig. 4*D*,*H*). The dependence on preparation time agrees with previous findings that goal-directed tuning of SLR gains requires a minimum of ∼300 ms of preparation (Rohlén et al., 2025).

**Figure 4.**
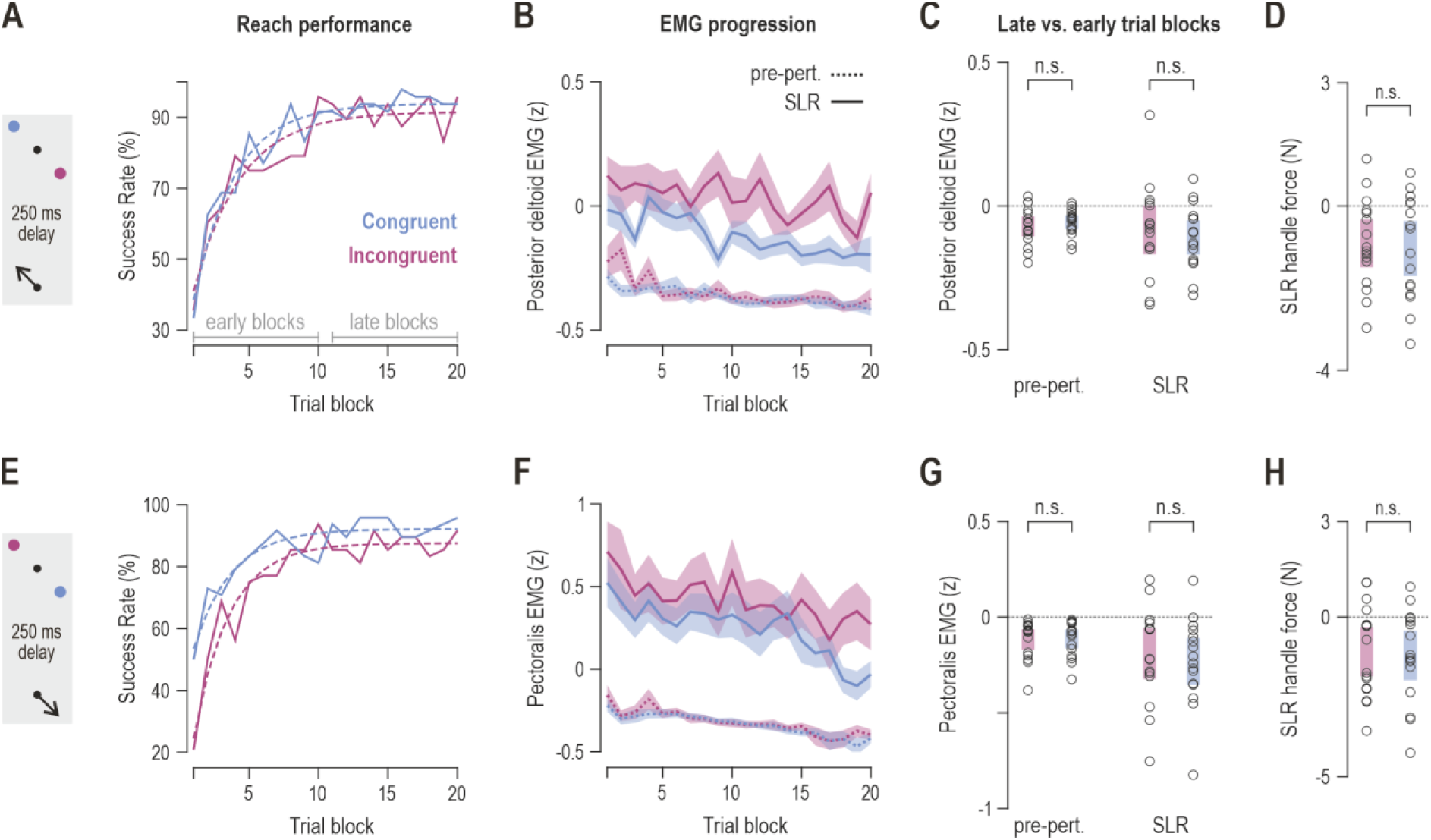
No goal-directed SLR tuning with a short preparatory delay. As Figure 3, but for trials with a short preparatory delay (250 ms). ***A–D***, Posterior deltoid; ***E–H***, pectoralis major. Despite comparable improvements in reaching performance (*A*, *E*), and similar gradual decline in pre-perturbation activity across target conditions (as in the long preparatory delay trials; Fig. 3), there were no significant goal-directed changes in SLR activity, pre-perturbation activity, or SLR-related handle force across the session. Single stars and n.s. as in Figure 3.

### Progressive SLR tuning across trial blocks

We next assessed whether SLR tuning developed progressively across consecutive trial blocks, beyond the difference between the early and late halves of the session. For each participant and target congruency, we computed the Spearman correlation coefficient (ρ) between EMG activity and trial block number (Fig. 5). Fisher z-transformed coefficients were compared between congruent and incongruent trials. To further control for background muscle activity, SLR correlations were also computed with pre-perturbation activity partialled out (within each load condition), and in the no-load condition alone, i.e., in the absence of any external load (see Materials and Methods).

**Figure 5.**
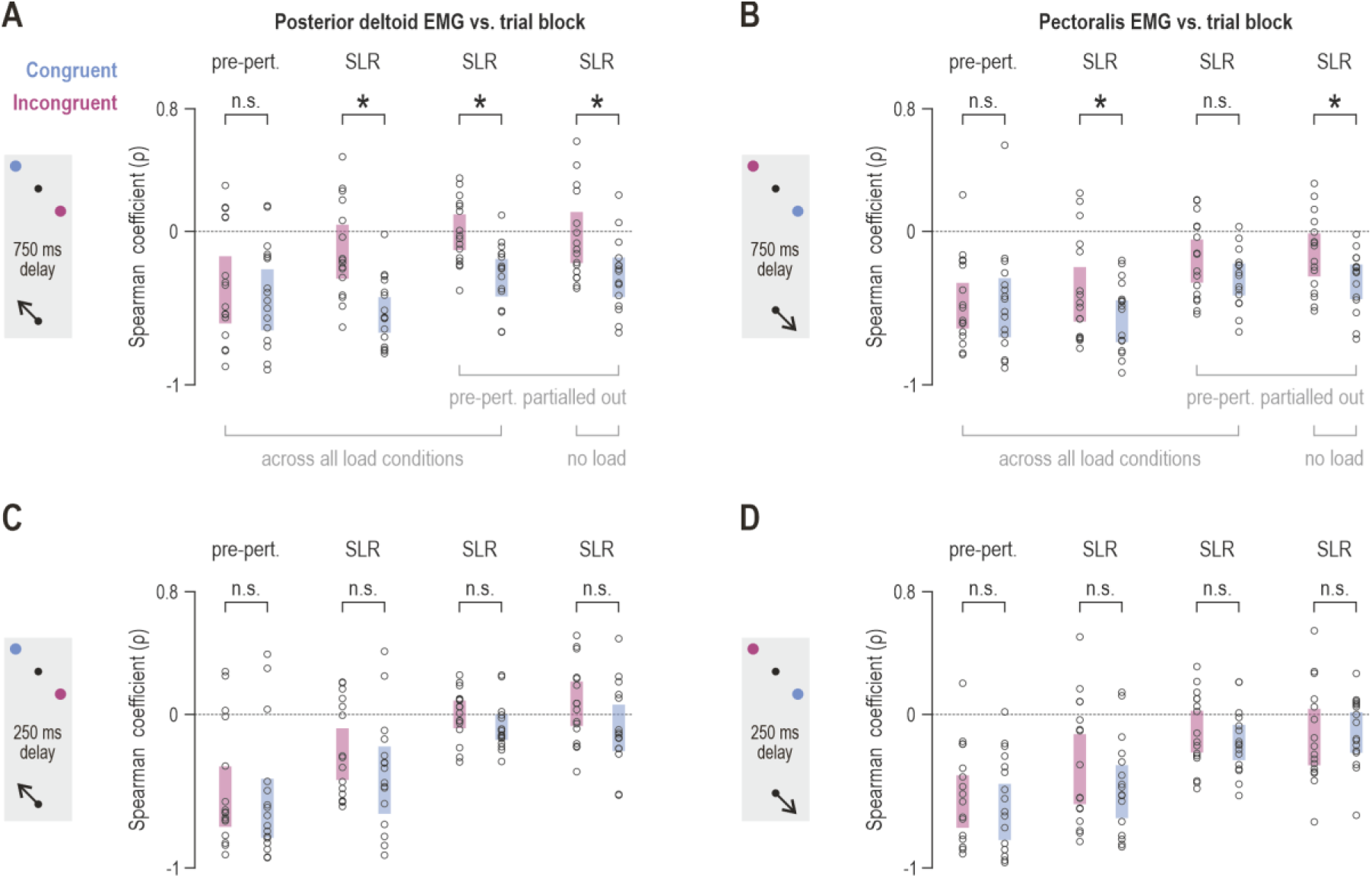
Progressive SLR tuning across trial blocks. ***A***, Spearman correlation coefficients (ρ) between posterior deltoid EMG activity and trial block for incongruent (purple) and congruent (blue) trials with a long preparatory delay (750 ms). From left to right: pre-perturbation activity and SLR activity across all load conditions; SLR activity with pre-perturbation activity partialled out (within each load condition), across all load conditions; and SLR activity with pre-perturbation activity partialled out in the no-load condition alone. Each circle represents a single participant, and shaded bars represent 95% confidence intervals (back-transformed from the Fisher z scale, on which statistical comparisons were performed). ***B***, As *A*, but for the pectoralis major. ***C–D***, As *A* and *B*, but for trials with a short preparatory delay (250 ms). Single stars indicate statistically significant paired comparisons of Fisher z-transformed coefficients at α = 0.05 (see Materials and Methods); n.s., not significant.

With the long preparatory delay, pre-perturbation activity showed no congruency-dependent progression in either muscle (posterior deltoid: Wilcoxon *W* = 66, *p* = 0.94; pectoralis: *t*(15) = −0.35, *p* = 0.73; Fig. 5*A*,*B*). In contrast, in the posterior deltoid, SLR activity was more negatively correlated with trial block in congruent than in incongruent trials (Fig. 5*A*; Wilcoxon *W* = 0, *p* = 3.1 × 10⁻⁵, r_rb_ = −1), and this remained the case when pre-perturbation activity was partialled out, both across all load conditions (*t*(15) = −4.32, *p* = 0.0006, *d*_z_ = −1.08) and in the no-load condition alone (*t*(15) = −2.5, *p* = 0.0247, *d*_z_ = −0.625). In the pectoralis, the progressive decrease in congruent relative to incongruent SLR responses was in the same direction (Fig. 5*B*; *t*(15) = −2.32, *p* = 0.035, *d*_z_ = −0.58). With pre-perturbation activity partialled out, it was weaker across all load conditions (*t*(15) = −1.93, *p* = 0.073), but reached significance again in the no-load condition alone (*t*(15) = −2.21, *p* = 0.043, *d*_z_ = −0.552), consistent with load-dependent gain-scaling restricting SLR tuning in this muscle. With the short preparatory delay, none of the pre-perturbation or SLR congruency comparisons was significant in either muscle (Fig. 5*C*,*D*; all *p* ≥ 0.1).

### LLR tuning across trial blocks

Finally, we examined whether the LLR (50–100 ms) progressed across the session in the same manner as the SLR (Fig. 6). In both muscles and at both preparatory delays, LLR responses across loads were markedly weaker when preparing to stretch rather than shorten the homonymous muscle throughout the session (Fig. 6*A*–*D*, left), with goal-directed tuning of the LLR already present in the first trial blocks. In addition, LLR responses in congruent trials decreased progressively across trial blocks. In the pectoralis, LLR responses in incongruent trials also decreased over the first few blocks with the long preparatory delay but remained relatively stable thereafter, whereas congruent responses continued to decrease across the session (Fig. 6*C*). Within participants, LLR activity was significantly more negatively correlated with trial block in congruent than in incongruent trials, in both muscles and at both preparatory delays (posterior deltoid, 750 ms: *t*(15) = −6.17, *p* = 1.8 × 10⁻⁵, *d*_z_ = −1.542; 250 ms: *t*(15) = −4.28, *p* = 0.00066, *d*_z_ = −1.07; pectoralis, 750 ms: *t*(15) = −6.44, *p* = 1.1 × 10⁻⁵, *d*_z_ = −1.61; 250 ms: *t*(15) = −4.1, *p* = 0.00095, *d*_z_ = −1.025; Fig. 6*A*–*D*, right). As expected, given the relative insensitivity of the LLR to automatic gain-scaling, partialling out pre-perturbation activity yielded similar results (posterior deltoid, 750 ms: *t*(15) = −5.95, *p* = 2.7 × 10⁻⁵, *d*_z_ = −1.487; 250 ms: *t*(15) = −5.14, *p* = 0.00012, *d*_z_ = −1.285; pectoralis, 750 ms: *t*(15) = −5.69, *p* = 4.3 × 10⁻⁵, *d*_z_ = −1.422; 250 ms: *t*(15) = −4.86, *p* = 0.00021, *d*_z_ = −1.215).

**Figure 6.**
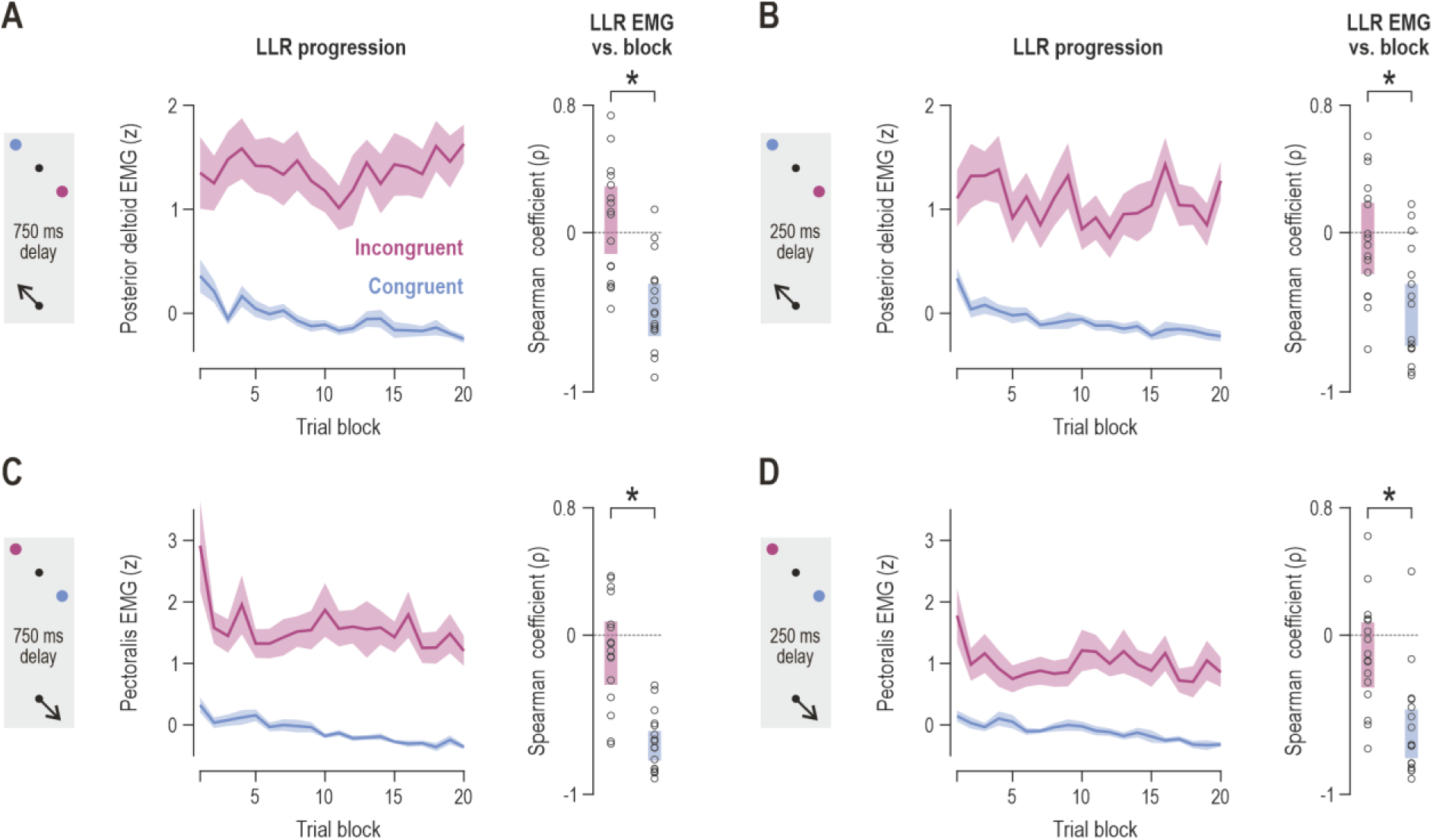
Progressive LLR tuning across trial blocks. ***A***, Posterior deltoid, long preparatory delay (750 ms). Left: mean z-scored LLR activity across trial blocks for incongruent (purple) and congruent (blue) trials; shading represents ±1 SEM. Right: Spearman correlation coefficients (ρ) between LLR activity and trial block (correlations with pre-perturbation activity partialled out are reported in the main text). Each circle represents a single participant, and shaded bars represent 95% confidence intervals, as in Figure 5. ***B***, As *A*, but with a short preparatory delay (250 ms). ***C–D***, As A and B, but for the pectoralis major. Single stars indicate statistically significant paired comparisons of Fisher z-transformed coefficients at α = 0.025 (Bonferroni correction across the two preparatory delays).

## Discussion

The central finding in this study is that spinal stretch reflexes can undergo rapid, experience-dependent tuning in the context of planned movement. We show that goal-directed suppression of the SLR can deepen progressively across a single session and weaken reflexive resistance to perturbations in the direction of the planned reach. Adaptive plasticity of spinal reflexes has classically been studied through operant conditioning, which induces an arbitrary change in reflex size that develops over days to weeks (Wolpaw et al., 1983; Thompson and Wolpaw, 2014). However, reflex gains are also adjustable on shorter timescales. Even in operant conditioning, a task-dependent adaptation of the H-reflex within the first few sessions is dissociable from the slower long-term change (Thompson et al., 2009). Moreover, the SLR adapts to recent movement history (Stratmann et al., 2024), and daily gait training progressively reshapes the H-reflex modulation pattern over days (Schneider and Capaday, 2003). Here, feedback was contingent on reaching performance rather than on reflex size, and the progressive change in SLR gains emerged as participants experienced the reaching task, with its direction dictated by the planned movement. Therefore, our results indicate that task constraints can shape the spinal SLR within a single session, challenging the view that flexible short-timescale feedback adaptation is the exclusive domain of transcortical (supraspinal) mechanisms.

Goal-dependent SLR suppression deepened over the session in both muscles (Fig. 3C,G), progressed across trial blocks within individual participants (Fig. 5A,B), and was accompanied by a corresponding decrease in reflex-related handle force (Fig. 3D,H). This suggests that reflex inhibition may facilitate movement by reducing antagonist resistance and limb stiffness during the executed reach. The gradual deepening of suppression from initially higher gains is consistent with optimal feedback control, in which feedback gains are continuously updated by the evolving motor plan: reflex gains track the formation of a decision and converge on task-specific values only as a goal is identified or becomes committed (Selen et al., 2012), and they are rapidly re-computed throughout a movement if goals suddenly change (Dimitriou et al., 2013). Within this framework, the goal-dependent SLR suppression we observe can be interpreted as the spinal-level expression of a feedback control policy that emerges once the upcoming reach has been specified. The directionality of this effect agrees with previous work showing categorical, goal-dependent modulation of muscle spindle afferents and stretch reflexes during movement preparation (Dimitriou, 2018; Papaioannou and Dimitriou, 2021; Torell et al., 2023a, b; Rohlén et al., 2025). The present study further shows that the magnitude of this goal-directed suppression evolves with experience within a single session.

Adaptive SLR tuning depended on having sufficient time to prepare for the upcoming reach. With a relatively long preparatory delay (750 ms), SLR tuning developed progressively across trial blocks (Figs. 3, 5), but no such tuning was detected when the preparatory delay was short (250 ms; Fig. 4). This dissociation supports the concept of advance specification of a feedback control policy (Ahmadi-Pajouh et al., 2012), in which the nervous system configures both the upcoming motor command and feedback gains before movement onset. Recent work has shown that ∼300 ms is typically sufficient for goal-directed tuning of SLR circuits in the context of planned reaching (Rohlén et al., 2025). Such tuning has also been previously shown with 750 ms and longer preparatory delays (Papaioannou and Dimitriou, 2021). A 750 ms delay is therefore not extreme and is arguably closer to everyday behavior than minimal-delay reaching; during naturalistic self-paced tasks, gaze typically shifts to an object ∼0.56 s before reach onset (Land et al., 1999), and prepared actions are often withheld further until the right moment. This timeline is also consistent with goal-directed fusimotor control, which relies on slowly conducting gamma efferents and electromechanical delays at the level of intrafusal muscle fibers (Dimitriou, 2022). The requirement for a minimal preparation time places the phenomenon within the framework of preparatory motor processing (Cohen et al., 2010; Haith and Bestmann, 2018), and provides one explanation for why earlier studies probing reflexes during ongoing, minimally prepared, or postural tasks may have missed such modulation. Reflex activity and its goal-directed modulation also appear distributed across much of the recording grid (Fig. 2B), and the denser spatial sampling of HDsEMG likely helped detect consistent SLR modulations. Such tuning may have gone undetected in earlier work relying on single-channel surface EMG.

Long-latency responses showed a large, categorical goal-directed modulation already in the initial trial blocks (Fig. 6), as widely known to be the case (Pruszynski and Scott, 2012; Scott, 2016). With experience, the LLR became progressively more inhibited when preparing to stretch the homonymous muscle, as did the SLR following the relatively long preparatory delay. The parallel trajectories of the two reflex components suggest that the SLR and LLR share a common feedback control policy that is updated incrementally with experience. This is in line with proposals that feedback gains are co-specified with the upcoming feedforward command (Ahmadi-Pajouh et al., 2012; Maeda et al., 2020), and with reports that long-latency feedback responses adapt across trials alongside reach adaptation (Cluff and Scott, 2013; Coltman and Gribble, 2020). Unlike the SLR, however, progressive LLR tuning was also evident with the short preparatory delay. As ∼250 ms is sufficient to adjust long-latency responses (Crevecoeur et al., 2020), this dissociation also argues against the SLR effects at the long preparatory delay reflecting an encroachment of long-latency activity into the SLR window, as this would be expected to produce comparable SLR effects at the short delay as well. It further suggests that the two reflex components may implement the same policy through partly different routes, with transcortical pathways reweighted faster (via access to central processing of incoming sensory information) than the slower tuning of the SLR.

The biomechanical context of the upcoming movement may also constrain the adaptive capacity of the SLR. The late-versus-early contrasts revealed equivalent goal-directed SLR tuning in both muscles, in both EMG and handle force (Fig. 3C,D,G,H). A difference between muscles emerged only at the finer, block-by-block resolution of the participant-level correlations: progressive tuning was significant in all comparisons for the posterior deltoid (Fig. 5A), whereas in the pectoralis it was significant across all loads but weaker once pre-perturbation activity was partialled out (p = 0.073), reaching significance again in the no-load condition (Fig. 5B). As the effect was consistently in the same direction, this pattern likely reflects a common effect in both muscles that was detected with less statistical power in the pectoralis. This could be because the pectoralis was more strongly affected by automatic gain-scaling, whereby reflex gain increases with background activation to maintain stability (Pruszynski et al., 2009; Pruszynski and Scott, 2012). Therefore, elevated baseline gain may not only enlarge the SLR but also reduce the range over which it can be tuned, to an extent that likely depends on how strongly the load engages each muscle. In addition to a top-down increase in α motor neuron excitability, this gain constraint may arise because independent control of muscle spindle sensitivity is co-opted by the tonically engaged homonymous muscle via α-γ co-activation (Vallbo, 1970; Dimitriou, 2014), β-motoneuron drive (Kakuda et al., 1998) and/or concomitant increases in intramuscular pressure (Torell and Dimitriou, 2024). These observations are consistent with a functional dissociation between postural stabilization, in which high spinal gains are advantageous, and goal-directed reaching, in which selective reduction of antagonist gain facilitates efficient execution (Weiler et al., 2021).

Background muscle activity declined slightly across the session, but this gradual decline was common across congruent and incongruent trials (Figs. 3B,F; 5A,B). In turn, this suggests a common, non-specific process such as fatigue or practice-related reduction of co-contraction (Osu et al., 2002). Through automatic gain-scaling (Pruszynski et al., 2009), a common decline in background activity would be expected to impact SLR responses equally in both congruencies. This decline appears, if anything, more pronounced following the short rather than the long preparatory delay (Figs. 4B,F vs. 3B,F; 5C,D vs. 5A,B). Yet SLR gains significantly diverged between congruent and incongruent trials following the relatively long preparatory delay only (Fig. 3C,G). Moreover, this divergence persisted within participants when pre-perturbation activity was partialled out, across all loads in the posterior deltoid, and in the no-load condition in both muscles (Fig. 5A,B), which in our task best approximates the context of reaching to grasp an object/load. Together, the above indicate that changes in α motor neuron output or automatic gain-scaling alone cannot account for the progressive goal-directed modulation of the SLR observed in this study.

Direct recordings of human muscle spindle activity during instructed-delay reaching have shown that primary (Ia) firing rates can decrease in a goal-dependent manner during preparation, in the absence of changes in EMG (Papaioannou and Dimitriou, 2021). In that study, however, participants had practiced the task before afferent recordings began. But our current findings raise the possibility that preparatory tuning of the sensory limb may itself be plastic within an experimental session. Such a mechanism is consistent with the broader proposal that human muscle spindles operate not as passive mechanoreceptors but as actively controlled, signal-processing devices (Dimitriou, 2022) whose sensitivity can be tailored to the task demands and shaped by learning (Dimitriou, 2016). It should be emphasized, however, that the SLR may be shaped at several points along the reflex arc. Presynaptic inhibition regulates transmission from Ia afferents to motoneurons (Eguibar et al., 1994; Lomelí et al., 1998), while neuromodulatory inputs control motoneuron excitability (Heckman et al., 2003), including through subthreshold changes not evident in surface EMG. Such mechanisms may well operate alongside independent fusimotor control, and the progressive SLR tuning observed here could reflect their combined effects.

In conclusion, the present study shows that spinal stretch reflexes can be progressively tuned with experience in a goal-dependent manner within a single session, driven by task constraints. This indicates that movement optimization relies not only on feedforward planning but also on proactive configuration of rapid feedback pathways, including spinal circuits. Together with the parallel progression of the LLR, our findings suggest that short- and long-latency feedback loops share a common, gradually updated control policy. The entire stretch reflex loop, including its fast spinal component, can therefore contribute to the rapidly progressing optimization of behavior.

## Acknowledgments

This work was supported by funds awarded to M.D. by the Strategic Research Fund of the Medical Faculty, Umeå University (FS 2.1.6-56-22) and Insamlingsstiftelserna (1025072). The authors would like to thank Anders Bäckström and Carola Hjältén for technical assistance.

## Supplementary Figures

**Figure S1.**
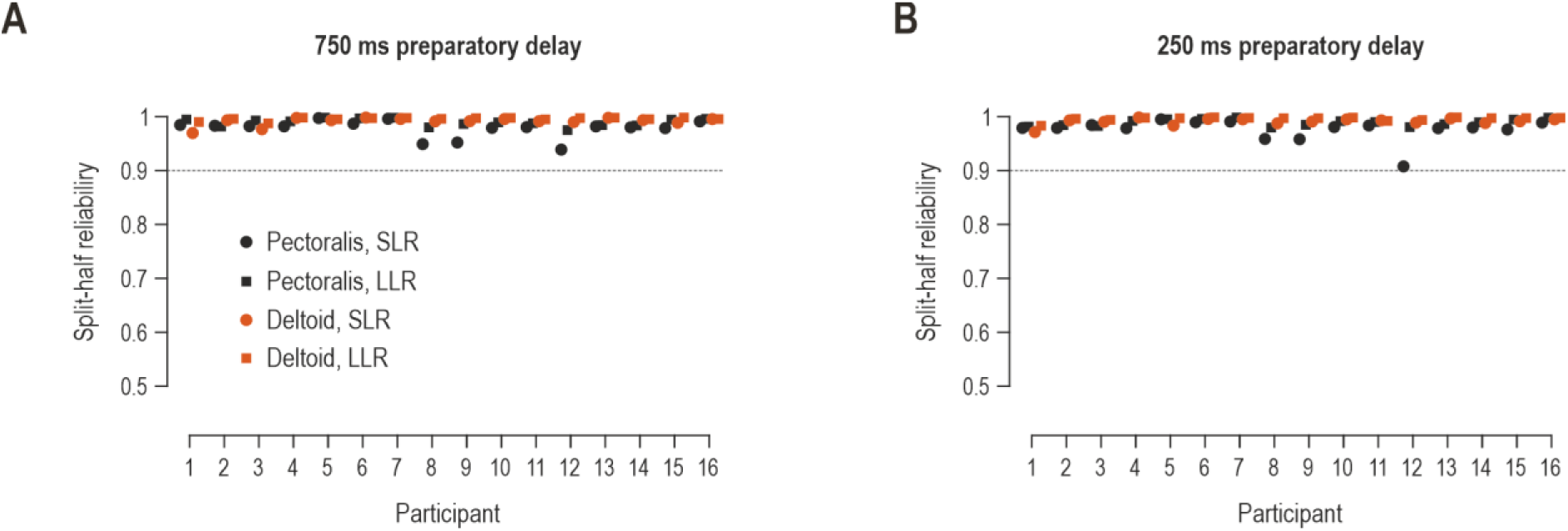
Split-half reliability of the aggregated EMG. ***A***, For trials with a long preparatory delay, reliability of the per- block EMG values across grid channels (Spearman–Brown corrected split-half correlation, within congruency; see Materials and Methods) for each participant, for the pectoralis (black) and posterior deltoid (orange). Circles represent the SLR epoch and squares the LLR epoch. The dotted line marks a reliability of 0.9. ***B***, As A, but for trials with a short preparatory delay.

**Figure S2.**
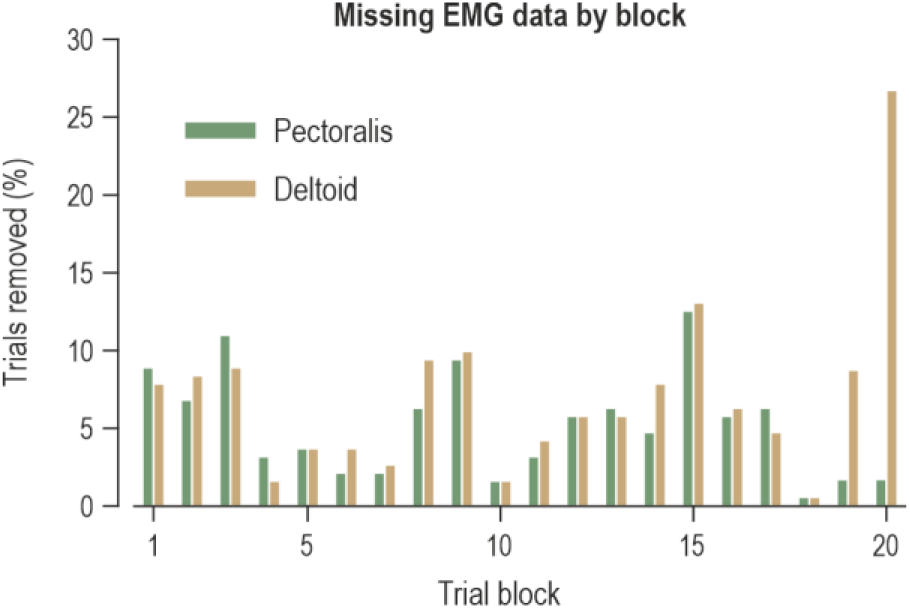
Distribution of missing EMG data. Percentage of trials removed as missing EMG data (including the contact-fault trials of one participant, block 20) in the analyzed conditions, for the pectoralis (green) and posterior deltoid (tan), by trial block.

**Figure S3.**
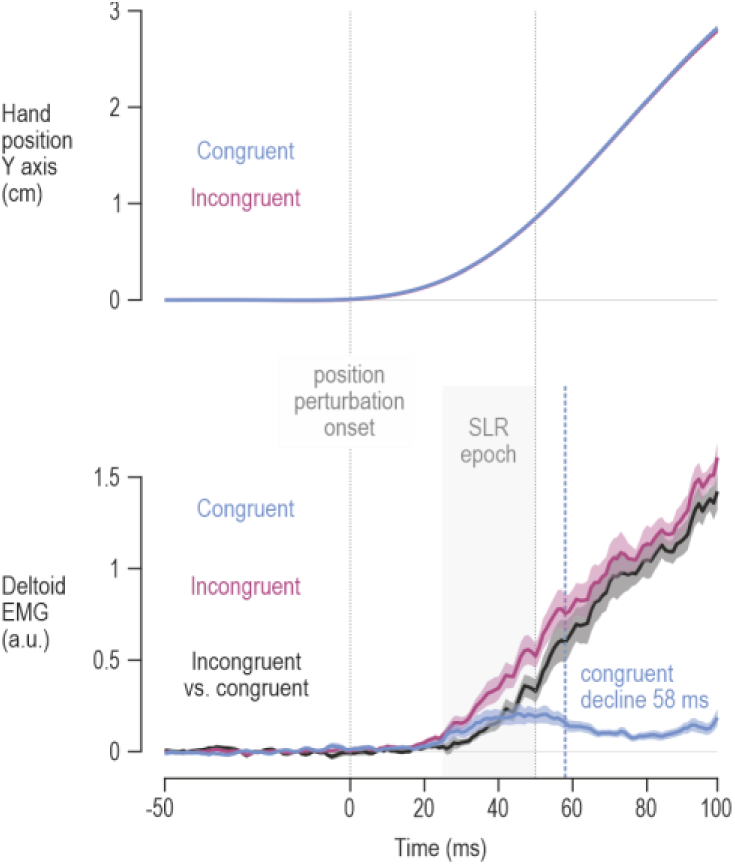
Onset of the SLR and LLR epochs. Data from the late trial blocks (11–20), pooled across load conditions, in which the posterior deltoid was stretched by the perturbation following a long preparatory delay (750 ms). Top panel: hand position along the task (Y) axis in congruent and incongruent trials (overlapping traces). Bottom panel: baseline-corrected deltoid EMG, normalized to each participant’s peak stretch response (a.u.), for congruent (blue) and incongruent (purple) trials, and their difference (incongruent − congruent; black). Vertical dotted lines mark perturbation onset (0 ms) and the SLR/LLR boundary (50 ms); the gray bar marks the SLR epoch (25–50 ms), and the blue dashed line the onset of the decline in congruent-trial EMG (58 ms; see Materials and Methods). Traces show the mean across participants (*n* = 16); shading represents ±1 SEM.

